# Low handgrip strength is closely associated with anemia among adults: A cross-sectional study using Korea National Health and Nutrition Examination Survey (KNHANES)

**DOI:** 10.1101/652545

**Authors:** Yu-mi Gi, Boyoung Jung, Koh-Woon Kim, Jae-Heung Cho, In-Hyuk Ha

**Author notes:** **Corresponding authors:** (BJ), (IHH).

## Abstract

**Background:** Anemia occurs because of insufficient hemoglobin, which provides oxygen to the body. Because of its close relationship with various illnesses, it must always be investigated clinically. Previous studies have demonstrated an association between hemoglobin concentration and handgrip strength. Thus, we aimed to analyze the association between handgrip strength and anemia in Korean adults to determine whether the handgrip strength test can be used as a tool to identify anemia.

**Methods:** The research subjects’ data were extracted from the 6^th^ and 7^th^ Korean National Health and Nutrition Examination Survey, between January 2013 and December 2017. Overall, data of 16,637 adults (weighted n= 9,734,598) were analyzed. Differences in sociodemographic factors (sex, age, education, income, and employment), lifestyle factors (alcohol consumption, smoking, and physical activity), and illness and health factors (body mass index, vitamin intake, iron intake, comorbid illnesses, and handgrip strength) by existence of anemia were analyzed using a chi square test. Binary logistic regression was performed to identify factors of anemia. Subgroup analysis, stratified by sex and age, was performed.

**Results:** Among Korean adults aged ≥ 19 years, 745,296 (7.7%) had anemia. Higher odds ratio (OR) of anemia occurred in the weak handgrip strength group compared to the strong handgrip strength group (OR=1.92, 95% CI: 1.58-2.33). The subgroup analysis showed a higher OR for anemia in the weak handgrip strength group than in the strong handgrip strength group, regardless of sex or age. However, the results showed that this association was greater for males (OR=2.13, 95% CI: 1.35-3.34) and for those aged ≥65 years (OR=1.92, 95% CI: 1.42-2.58).

**Conclusion:** This study showed a strong association between handgrip strength and anemia, which was particularly strong for males and those aged ≥65 years. Therefore, it is anticipated that handgrip strength can be used in anemia screening tests as a useful tool.

## Introduction

Anemia occurs because of insufficient hemoglobin, which provides oxygen to the body. It is one of the major clinical problems affecting about 12% of the elderly [1]. The World Health Organization (WHO) defines anemia as hemoglobin values below 12 g/dl for females and 13 g/dl for males [2]. Anemia affects mortality rate [3], disability [4], the decline in physical activity level [4, 5], and decline in the quality of life [6] in the elderly. Furthermore, since anemia has a close relationship with many illnesses, it is recommended that when hemoglobin level is lower than the normal, it should always be followed-up clinically [5]. Hence, the prevalence of anemia can then be reduced by appropriate treatment or by preventing the underlying illnesses that are related to aging [7].

Muscle weakness is related to poor physical activity and incident mobility limitations in the elderly [3, 6, 8–11]. Handgrip strength, a simple, cost-effective tool used to measure strength, is utilized in the diagnosis of muscle weakness in epidemiological studies [10]. Handgrip strength is also a non-invasive and cheap tool that can help in identifying those with a higher likelihood of anemia, and can be used as a tool to identify anemia in the community [12]. Some research has reported that handgrip strength in middle-age (45-65 years) is related to functional limitations and disabilities at least 25 years later [3, 11, 13]. Therefore, handgrip strength has been considered a factor predicting healthy aging [11]. Handgrip strength is also related to the strength of other muscle groups, and therefore can be a good index to represent overall strength [14]. Therefore, low handgrip strength signifies low muscle mass and poor clinical mobility [15].

Oxidative stress is one of the most important mechanism to explain the occurrence of age-related illnesses [16] including the decline in strength [17]. Because hemoglobin concentration plays an important role in the oxidation of blood [18, 19], anemia, defined by hemoglobin, represents a decline in the oxygenation function and can therefore be related to low strength [4].

Aging is the most frequent cause of reduced muscle strength [20], and anemia is commonly related to aging [21]. Previous research in Australian elderly found a direct relationship between reduction in hemoglobin level and the reduction in handgrip strength [22]. In addition, in a research among elderly participants aged ≥65 years, anemia patients, defined by the hemoglobin level, had significantly lower handgrip strength compared to patients without anemia [4]. From these previous studies, it is evident that the hemoglobin level can be directly related to handgrip strength and that this is especially pronounced in the elderly population; however, the mechanism underlying this is unclear. Moreover, research on the association between handgrip strength and anemia in Korean adults has provided insufficient data for a clear conclusion. Therefore, in the present study, we aimed to determine whether an association exists between handgrip strength and anemia based on the data from the 6^th^ and 7^th^ Korean National Health and Nutrition Examination Survey (KNHANES).

## Materials and Methods

### Study design and population

The present study used the data from the 6^th^ and 7^th^ KNHANES that was conducted from January 2013 to December 2017 [17]. KNHANES is a 3-yearly cross-sectional research, which includes examinations, health surveys, and nutritional surveys, and involves a representative sample of the entire Korean population, using a stratified cluster-sampling method.

The KNHANES study sample consisted of Korea citizens (residents in group facilities, such as nursing homes, military, and prison as well as foreigners are excluded). To increase the representativeness of the sample and the accuracy of assumptions, sampling was designed to distinguish between province and city, neighborhood, towns, and townships, and the types of home, to extract sampling regions (investigated district). We used the data from 2013-2017 to measure handgrip strength. The subject exclusion criteria were as follows: missing data on anemia (n=8,928) and on handgrip strength (n=7,556), those aged under 19 years (n=2,307), and other missing data (n=3,797). Ultimately, the analyses were performed using data of 16,637 subjects. Final analysis set considering weight 9,734,598 subjects (Fig 1).

**Fig 1.**
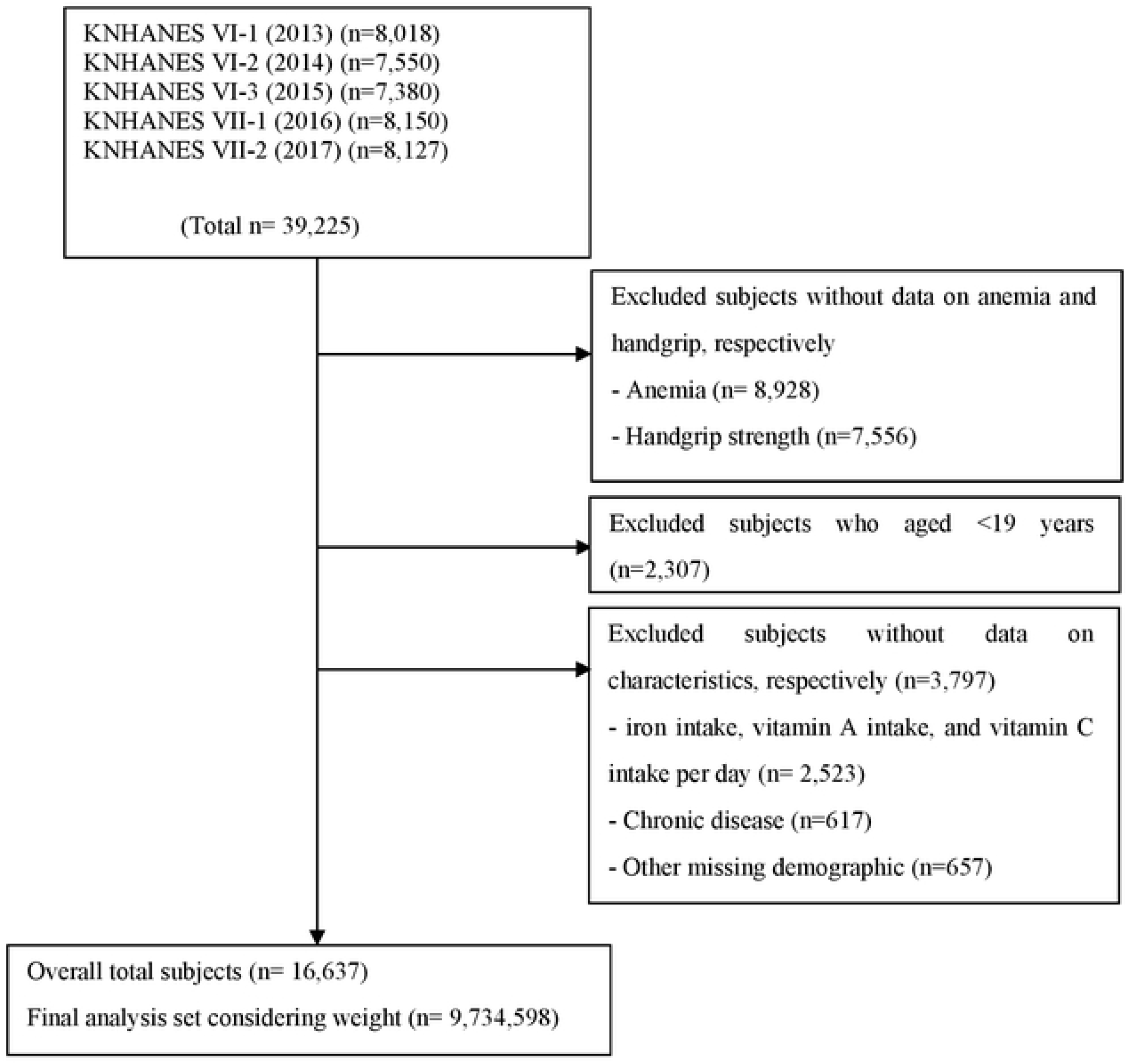
Subjects’ flow diagram.

### Outcomes and other variables

#### Measurement of handgrip strength

Handgrip strength, an indicator of muscle strength, was evaluated using a handgrip strength test according to the recommendations of the Institute of Medicine [14]. For the standard value of handgrip strength, we used the standard for the skeletal muscle mass (26 kg for men, 16 kg for women) suggested by the North American Foundation for the National Institutes of Health Sarcopenia Project published in 2015 [22]. The handgrip strength for each hand was measured three times using a digital grip strength dynamometer (TKK 5401; Takei Scientific Instruments Co, Ltd, Tokyo, Japan). Trained medical technicians instructed the study subjects while in a sitting position to hold the dynamometer with the distal interphalangeal finger joints at a 90° angle to the handle, and to squeeze the handle as firmly as they could. Another handgrip strength measurement was done during expiration, after the subject had slowly stood up. Study subjects conducted 3 attempts per hand, with a 1-minute rest period between each attempt to reduce the effect of fatigue due to repetition. The value for the handgrip strength measurements was the average of the three measurements, for either hand [23]. Using this mean handgrip strength values, subjects were categorized into the weak (<26 kg for men, <16 kg for women) and strong (≥26 kg for men, ≥16 kg for women) groups.

#### Measurement of hemoglobin (Hb) levels

Hemoglobin (Hb) was used as an index to determine anemia. In accordance with the WHO standards, male and female subjects with Hb less than 13 and 12 g/dL, respectively, were defined as anemia patients [13].

#### Description of demographic and other characteristics of the study population

We analyzed the general characteristics, socioeconomic background characteristics, and lifestyle habits of the patients. Body mass index (BMI), a value derived by dividing the weight with the square of height, was categorized into <18.5 kg/m^2^ (underweight), 18.5-24.9 kg/m^2^ (normal), and ≥25.0 kg/m^2^ (overweight) [16]. For smoking, subjects were categorized into nonsmoker, ex-smoker, or current smoker. The frequency of alcohol consumption was divided into four categories (‘none,’ ‘less than 1 time per month’, ‘1-4 times per month’, and ‘more than 5 times per month.’) For employment, subjects were categorized into ‘unemployed (student, housewife, etc.)’ or ‘employed.’ For marital status, subjects were categorized into three groups (’married-cohabiting’ for those who had spouses or cohabiting partners; ‘married-non cohabiting, bereaved, or divorced’ for those who were living apart, bereaved, divorced, etc.; or ‘unmarried’ for those who were unmarried). Household income was categorized into four groups (low, low-moderate, moderate-high, and high). Educational level was also categorized into four groups (elementary school or less, middle school, high school, and college or over). Level of daily workout refers to the frequency of strength exercise per week, and was categorized into four groups (’none’; ‘light’ [1 day]; ‘moderate’ [2∼3 days]; and ‘heavy/extreme’ [4 days or more]).

#### Description of nutritional characteristics and comorbidity in the study population

For nutritional characteristics, iron intake, vitamin A intake, and vitamin C intake per day were categorized as ‘insufficient’ if the intake was lower than the recommended level by sex and age and ‘sufficient’ if the intake was above the recommended level. All blood samples were drawn after fasting and were analyzed within 24 hours of being drawn. The 2015 standards of the Dietary Reference Intakes for Koreans (KDRIs) were used for iron and vitamin C intake per day. For vitamin A intake, the 2010 KDRIs standards [24] were used with identical units (μg RE/day) for KHANES and KDRIs [25] (μg RAE/day), in 2015. Vitamin A intake per day for males and females aged 19-49 and ≥50 years were 750 and 700 μg RE/day; and 650 and 600 μg RE/day, respectively. Vitamin C intake per day was 100 mg for both males and females over 19 years old. Iron intake per day for males were 10 and 9 mg/day for those aged 19-64 and ≥65 years, respectively; while for females aged 19-49, 50-74, and ≥75 years, these were 14, 8, and 7 mg/day, respectively.

For comorbid diseases, hypertension, cancer, hyperlipidemia, cardiovascular disease, kidney disease, bone disease, and thyroid disease were included. Subjects were categorized into a ‘yes’ or ‘no’ group based on the diagnosis or absence of at least one of these diseases by a doctor.

### Statistical analysis

Because KNHANES data were obtained from a complex sampling design, all statistical analyses methods used complex sample analyses considering weight, stratified variables, and cluster sampling. Differences in sociodemographic factors (sex, age, education, income, and employment), lifestyle factors (alcohol consumption, smoking, and physical activity), and illness and health factors (BMI, vitamin intake, iron intake, comorbid illnesses, and handgrip strength) due to presence of anemia were analyzed using a complex sample analysis of chi square (*χ*^2^) test. Categorical variables were presented as proportions (n, %) using a *χ*^2^ test of independence, while continuous variables were presented with estimated value ± standard error (SE) using the general linear model.

Chi-square test was performed on risk factors associated with anemia. Furthermore, to identify factors associated with anemia, odds ratio (OR) and 95% confidence intervals (95% CI) were obtained through a binary logistic regression analysis after setting anemia as the dependent variable and handgrip strength as the main independent variable, while controlling for sociodemographic factors, lifestyle habit factors, and disease-related and health factors. The association between handgrip strength and anemia were subgroup-analyzed by sex (male/female) and age (younger or older than 65). All statistical analyses were performed using SPSS version 25.0 (SPSS Inc., Chicago, IL, USA) and SAS version 9.4 (SAS Institute Inc, Cary, NC), and the significance level of all tests were defined as p-value less than 0.05.

### Ethics statement

The KNHANES 6^th^ and 7^th^ were conducted by the Korea Center for Disease Control and Prevention (KCDC). All survey protocols were approved by the institutional review board (IRB) of the KCDC (approval numbers: 2013-07CON-03-4C, 2013-12EXP-03-5C, and 2015-01-02-6C). Informed consent was obtained from all participants when the surveys were conducted. Original data are publicly available for free in the KNHANES website (http://knhanes.cdc.go.kr) for purposes such as academic research. Approval of IRB was not required because the study did not deal with any sensitive information, but rather accessed only publicly available data from the KNHANES (JASENG IRB File No. 2019-03-009).

## Result

Table 1 shows the general characteristics of the subjects. Patients with anemia were 7.7% overall. Majority of subjects (51.2%) were female, of which, 11.9% had anemia. In males only 3.2% had anemia. Overall, 6.7% of subjects and 15.8% were in the strong and weak handgrip strength groups, respectively (p<0.0001). The mean age of subjects with anemia was higher (50.27 years) than for those in the group without anemia (46.14 years) (p<0.0001). By marital status, those in the married-not cohabiting, bereaved, or divorced group had a higher percentage (11.6%) of anemia than those in the married-cohabiting (8.1%) and the unmarried (4.7%) groups (p<0.0001). The prevalence rates of anemia in the unemployed and employed groups were 10% and 6.3%, respectively (p<0.0001).

**Table 1.**
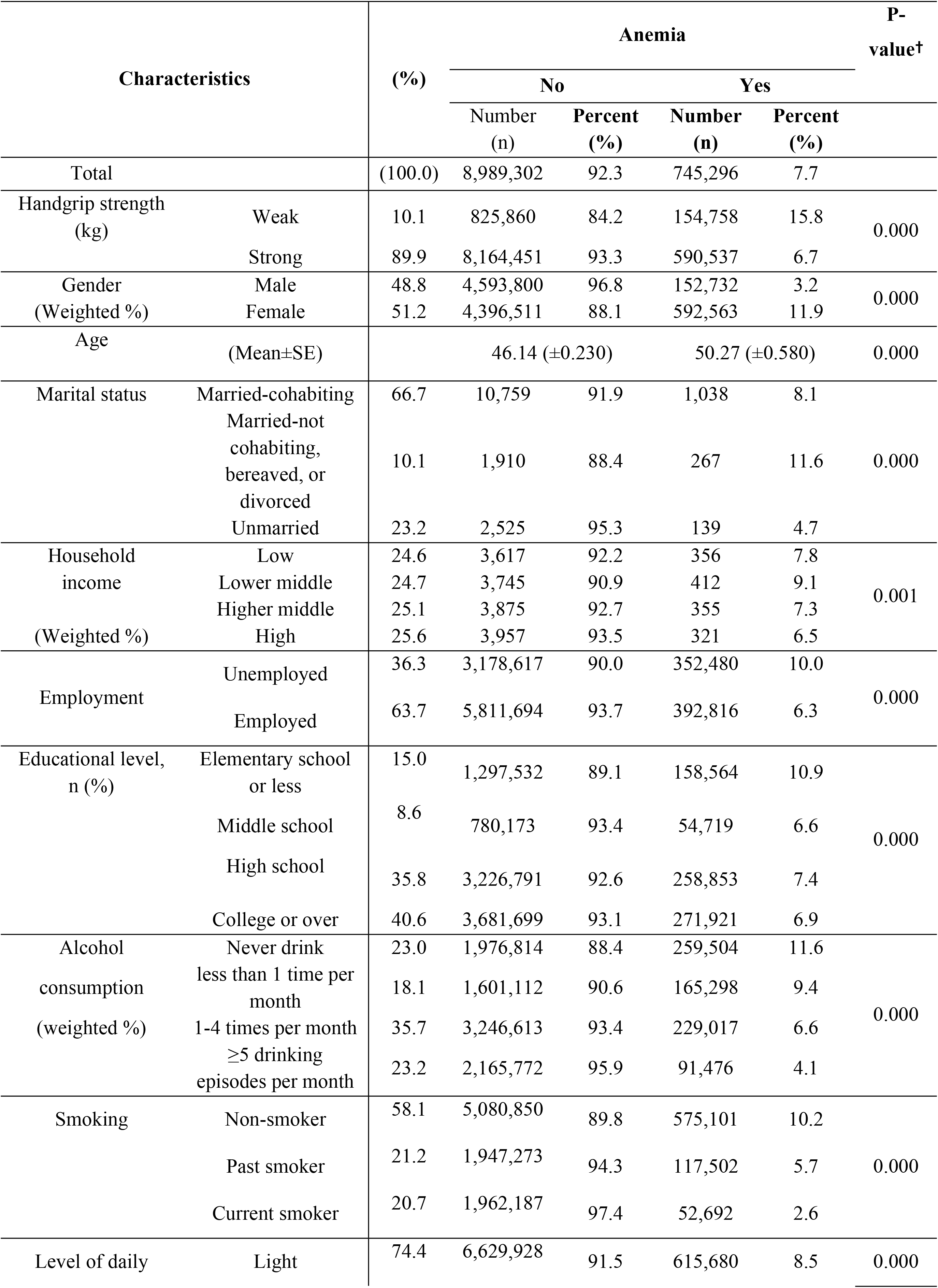
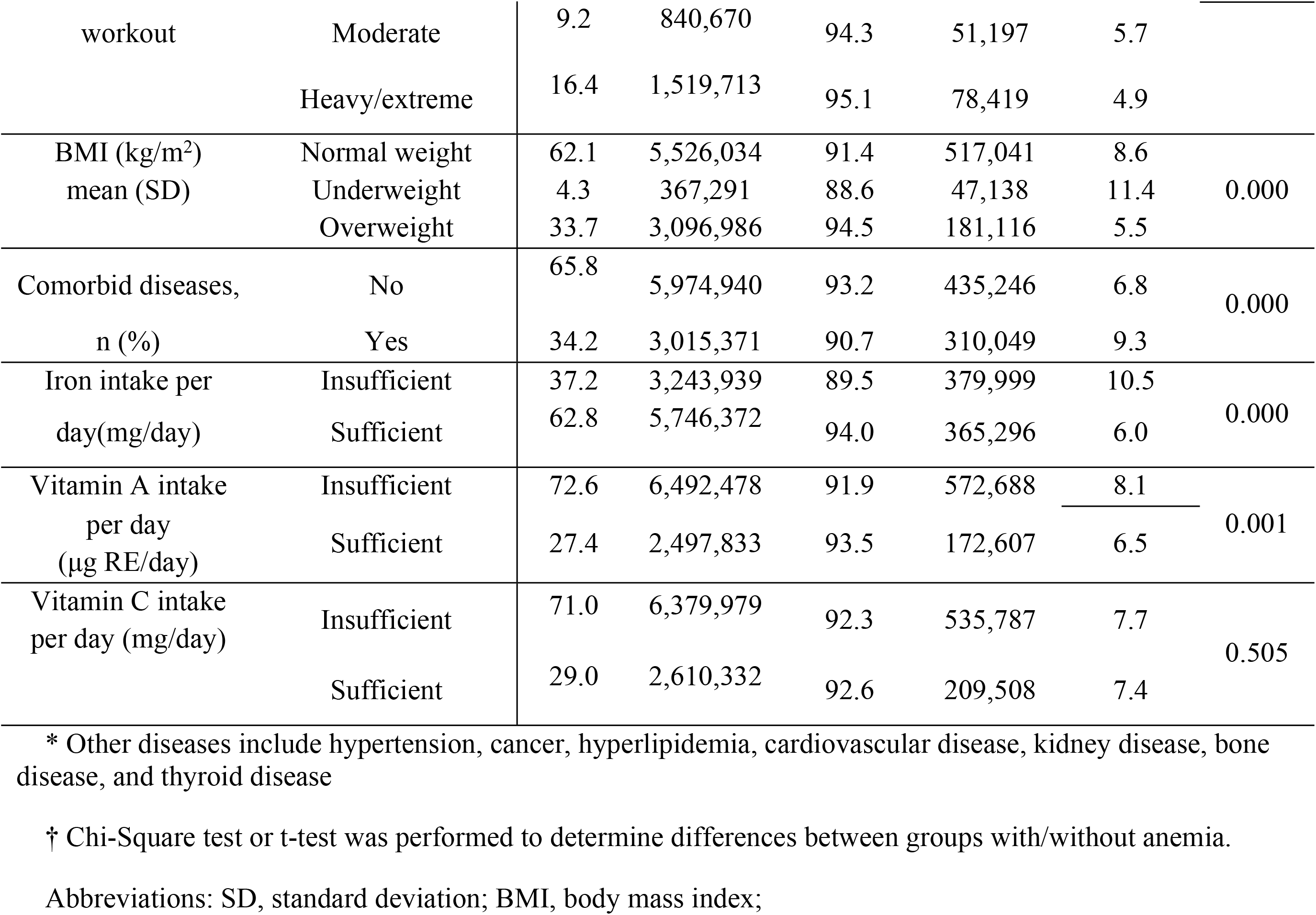
Characteristics of the study population.

By individual lifestyle habit factors, by the level of daily workout, in the light (1 day) daily workout group 615,680 (8.5%) subjects had anemia, and had higher prevalence of anemia than those in the moderate (5.7%) and heavy/extreme (4.9%) daily workout groups, (p<0.0001). The prevalence rate of anemia in the underweight group (11.4%) was higher than that in the normal weight or overweight groups (8.6% and 5.5%, respectively) (p<0.0001).

For the nutritional and disease-related factors, the prevalence rate of anemia in the group with comorbid diseases was 2.5% higher than in the group without (p<0.0001). For iron intake, the prevalence rate of anemia was 3.5% greater in the insufficient group compared to the sufficient group (p<0.0001). For vitamin A intake, the prevalence rates of anemia was also higher (8.1%) for the insufficient group than for the sufficient group (6.5%) (p=0.001). However, no difference between groups was found in vitamin C intake per day (p=0.505).

Table 2 shows the characteristic of subjects by sex. For handgrip strength, in both males (17.0% vs. 3.1%) and females (16.5% vs. 10.7%) the prevalence rate was significantly higher in the weak group than in the strong group (p<0.0001). Only in males were the mean ages of persons with and without anemia with a difference of about 18 years. Similarly, anemia prevalence rate was only higher in the underweight male group (15.6%) compared to the normal (5.8%) and overweight (3.1%) groups (p<0.0001), unlike for females (p=0.074). Anemia prevalence rate was also higher only in males with comorbid diseases (9.4%) than for those without comorbid disease (2.0%) (p<0.0001); but not for females, although the group without comorbid disease (11.8%) had slightly higher prevalence rate than those with comorbid diseases (10.9%) (p=0.065). The prevalence rate of anemia was lower in the iron intake sufficient groups (4.8% for males and 9.7% for females) compared to the insufficient groups (5.7% for males and 13.6% for females), but this was not significant in males (p=0.492).

**Table 2.** Characteristics of the study population by anemia according to sex.

|  |  | Male |  |  |  |  |  | Female |  |  |  |  |  |
| --- | --- | --- | --- | --- | --- | --- | --- | --- | --- | --- | --- | --- | --- |
|  |  | Total(n) | anemia |  |  |  | P-value† | Total(n) | anemia |  |  |  | P-value† |
|  |  |  | No |  | Yes |  |  |  | No |  | Yes |  |  |
|  |  |  | Number<br>(n) | Percent<br>(%) | Number<br>(n) | Percent<br>(%) |  |  | Number<br>(n) | Percent<br>(%) | Number<br>(n) | Percent<br>(%) |  |
| Total |  | 4746532 | 4593800 | 96.8 | 152732 | 3.2 |  | 4989075 | 4396511 | 88.1 | 592563 | 11.9 |  |
| Hand grip(kg)-2Q | weak | 436455 | 378554 | 86.7 | 57901 | 13.3 | 0.000 | 544164 | 447307 | 82.2 | 96858 | 17.8 | 0.000 |
|  | strong | 4310077 | 4215246 | 97.8 | 94831 | 2.2 |  | 4444910 | 3949205 | 88.8 | 495706 | 11.2 |  |
| Age(yr) | (Mean±SE) |  | 45.25±0.278 |  | 63.62±0.914 |  | 0.000 |  | 47.07±0.260 |  | 46.82±0.609 |  | 0.000 |
|  |  |  | 45.84±0.279 |  |  |  |  |  | 47.04±0.252 |  |  |  |  |
| Region | Urban area | 3971297 | 3848194 | 96.9 | 123103 | 3.1 | 0.192 | 4263914 | 3762553 | 88.2 | 501361 | 11.8 | 0.503 |
|  | Rural area | 775235 | 745606 | 96.2 | 29629 | 3.8 |  | 725161 | 633959 | 87.4 | 91202 | 12.6 |  |
| Marital status | Married-cohabiting | 3160785 | 3038119 | 96.1 | 122666 | 3.9 | 0.000 | 3330775 | 2929202 | 87.9 | 401573 | 12.1 | 0.146 |
|  | Married-not cohabiting,<br>bereaved, or divorced | 240072 | 222432 | 92.7 | 17640 | 7.3 |  | 745488 | 648825 | 87.0 | 96663 | 13.0 |  |
|  | Unmarried | 1345675 | 1333249 | 99.1 | 12426 | 0.9 |  | 912811 | 818485 | 89.7 | 94327 | 10.3 |  |
| Income | Low | 1180706 | 1135023 | 96.1 | 45682 | 3.9 | 0.115 | 1216663 | 1075659 | 88.4 | 141005 | 11.6 | 0.003 |
|  | Lower middle | 1160131 | 1120798 | 96.6 | 39334 | 3.4 |  | 1242594 | 1064273 | 85.6 | 178321 | 14.4 |  |
|  | Higher middle | 1190316 | 1153349 | 96.9 | 36967 | 3.1 |  | 1255929 | 1114039 | 88.7 | 141889 | 11.3 |  |
|  | High | 1215379 | 1184630 | 97.5 | 30749 | 2.5 |  | 1273889 | 1142541 | 89.7 | 131349 | 10.3 |  |
| Employment | Unemployed | 1162602 | 1091803 | 93.9 | 70799 | 6.1 | 0.000 | 2368495 | 2086815 | 88.1 | 281680 | 11.9 | 0.972 |
|  | Employed | 3583930 | 3501997 | 97.7 | 81933 | 2.3 |  | 2620580 | 2309697 | 88.1 | 310883 | 11.9 |  |
| Education Level | Elementary school or less | 504239 | 454972 | 90.2 | 49267 | 9.8 | 0.000 | 951858 | 842561 | 88.5 | 109297 | 11.5 | 0.014 |
|  | Middle school | 403036 | 384138 | 95.3 | 18898 | 4.7 |  | 431856 | 396034 | 91.7 | 35822 | 8.3 |  |
|  | High school | 1770921 | 1718534 | 97.0 | 52387 | 3.0 |  | 1714723 | 1508257 | 88.0 | 206465 | 12.0 |  |
|  | College or over | 2066023 | 2034615 | 98.5 | 31408 | 1.5 |  | 1887597 | 1647084 | 87.3 | 240513 | 12.7 |  |
| Alcohol consumption | Never drink | 700037 | 644773 | 92.1 | 55264 | 7.9 | 0.000 | 1536281 | 1332041 | 86.7 | 204240 | 13.3 | 0.011 |
|  | less than 1 time per month | 563460 | 545632 | 96.8 | 17828 | 3.2 |  | 1202949 | 1055480 | 87.7 | 147469 | 12.3 |  |
|  | 1-4 times per month | 1824263 | 1781506 | 97.7 | 42758 | 2.3 |  | 1651367 | 1465107 | 88.7 | 186260 | 11.3 |  |
|  | ≥5 times per month | 1658772 | 1621890 | 97.8 | 36882 | 2.2 |  | 598477 | 543883 | 90.9 | 54594 | 9.1 |  |
| Smoking | Non-smoker | 1244812 | 1208848 | 97.1 | 35964 | 2.9 | 0.000 | 4411140 | 3872002 | 87.8 | 539137 | 12.2 | 0.029 |
|  | Past smoker | 1766677 | 1680389 | 95.1 | 86287 | 4.9 |  | 298099 | 266884 | 89.5 | 31215 | 10.5 |  |
|  | Current smoker | 1735043 | 1704562 | 98.2 | 30481 | 1.8 |  | 279836 | 257625 | 92.1 | 22211 | 7.9 |  |
| Level of daily workout | Light | 3142034 | 3024438 | 96.3 | 117595 | 3.7 | 0.000 | 4103574 | 3605490 | 87.9 | 498084 | 12.1 | 0.143 |
|  | Moderate | 531443 | 524745 | 98.7 | 6697 | 1.3 |  | 360424 | 315925 | 87.7 | 44499 | 12.3 |  |
|  | Heavy/extreme | 1073055 | 1044616 | 97.3 | 28439 | 2.7 |  | 525076 | 475097 | 90.5 | 49980 | 9.5 |  |
| Stress level | High | 1205557 | 1183127 | 98.1 | 22430 | 1.9 | 0.000 | 1459232 | 1283937 | 88.0 | 175295 | 12.0 | 0.982 |
|  | Middle | 2777411 | 2688397 | 96.8 | 89013 | 3.2 |  | 2801742 | 2470974 | 88.2 | 330768 | 11.8 |  |
|  | Low | 746842 | 706690 | 94.6 | 40152 | 5.4 |  | 705094 | 620951 | 88.1 | 84143 | 11.9 |  |
|  | Zero | 16722 | 15585 | 93.2 | 1136 | 6.8 |  | 23007 | 20649 | 89.8 | 2357 | 10.2 |  |
| BMI (kg/m2) | Normal weight | 2725568 | 2623997 | 96.3 | 101571 | 3.7 | 0.000 | 3317508 | 2902038 | 87.5 | 415470 | 12.5 | 0.074 |
|  | Underweight | 125363 | 112066 | 89.4 | 13297 | 10.6 |  | 289066 | 255225 | 88.3 | 33842 | 11.7 |  |
|  | Overweight | 1895601 | 1857737 | 98.0 | 37864 | 2.0 |  | 1382501 | 1239249 | 89.6 | 143252 | 10.4 |  |
| other diseases | No | 3242557 | 3200355 | 98.7 | 42201 | 1.3 | 0.000 | 3167629 | 2774584 | 87.6 | 393045 | 12.4 | 0.065 |
|  | Yes | 1503975 | 1393444 | 92.7 | 110531 | 7.3 |  | 1821445 | 1621927 | 89.0 | 199519 | 11.0 |  |
| Iron intake per day(mg/day) | insufficient | 1117883 | 1079199 | 96.5 | 38684 | 3.5 | 0.492 | 2506055 | 2164739 | 86.4 | 341315 | 13.6 | 0.000 |
|  | sufficient | 3628649 | 3514600 | 96.9 | 114048 | 3.1 |  | 2483020 | 2231772 | 89.9 | 251248 | 10.1 |  |
| Vitamin A intake per day (µg RE/day) | insufficient | 3362075 | 3243391 | 96.5 | 118685 | 3.5 | 0.029 | 3703091 | 3249087 | 87.7 | 454004 | 12.3 | 0.087 |
|  | sufficient | 1384457 | 1350409 | 97.5 | 34047 | 2.5 |  | 1285984 | 1147424 | 89.2 | 138560 | 10.8 |  |
| Vitamin C intake per day (mg/day) | insufficient | 3444984 | 3338160 | 96.9 | 106824 | 3.1 | 0.357 | 3470782 | 3041819 | 87.6 | 428963 | 12.4 | 0.048 |
|  | sufficient | 1301548 | 1255640 | 96.5 | 45908 | 3.5 |  | 1518293 | 1354692 | 89.2 | 163601 | 10.8 |  |

\* Other diseases include hypertension, cancer, hyperlipidemia, cardiovascular disease, kidney disease, bone disease, and thyroid disease
† Chi-Square test or t-test was performed to determine differences between groups with/without anemia according to sex.
Abbreviations: yr, year; SE, standard error; BMI, body mass index;

Table 3 shows the characteristics of subjects by age and anemia status. The mean age of subjects aged <65 and ≥65 years were 41.72 and 72.41 years, respectively (p<0.0001). In the <65 and ≥65 years, respectively, anemia prevalence rate was higher in the weak than the strong handgrip strength group (<65 years, 10.9 vs. 6.5%, p<0.0001) and (≥65 years, 20.2% vs. 8.7%, p<0.0001), respectively. For the <65-year-olds, more female than males had anemia (11.6% vs. 1.7%) and a slight difference for the ≥65-year-olds (13.1% vs. 12.0%), respectively. Significant (<65 years, 8.7% vs. 5.9%, p<0.0001) and non-significant (≥65 years, p=0.207) difference occurred in the prevalence rate of anemia in the unemployed and employed groups, respectively. For alcohol consumption, the prevalence rate of anemia was lower for those who consume alcohol and especially those with a higher frequency of alcohol consumption, regardless of age (p<0.0001). For the level of daily workout, the prevalence rate of anemia was higher in the light group compared to the moderate or heavy/extreme group for all subjects regardless of age (p<0.0001). Also for BMI, the prevalence rate of anemia in the underweight group (<65 years, 10.1%; ≥65 years, 23.2%) was far higher than the prevalence rate among the normal weight or overweight groups (p<0.0001), regardless of age. Furthermore, for iron intake, the prevalence rate of anemia in the insufficient group (<65 years 9.7%; ≥65 years 15.7%), was higher than the prevalence rate in the sufficient group (<65 years 4.9%; ≥65 years, 11.3%) for both age groups.

**Table 3.**
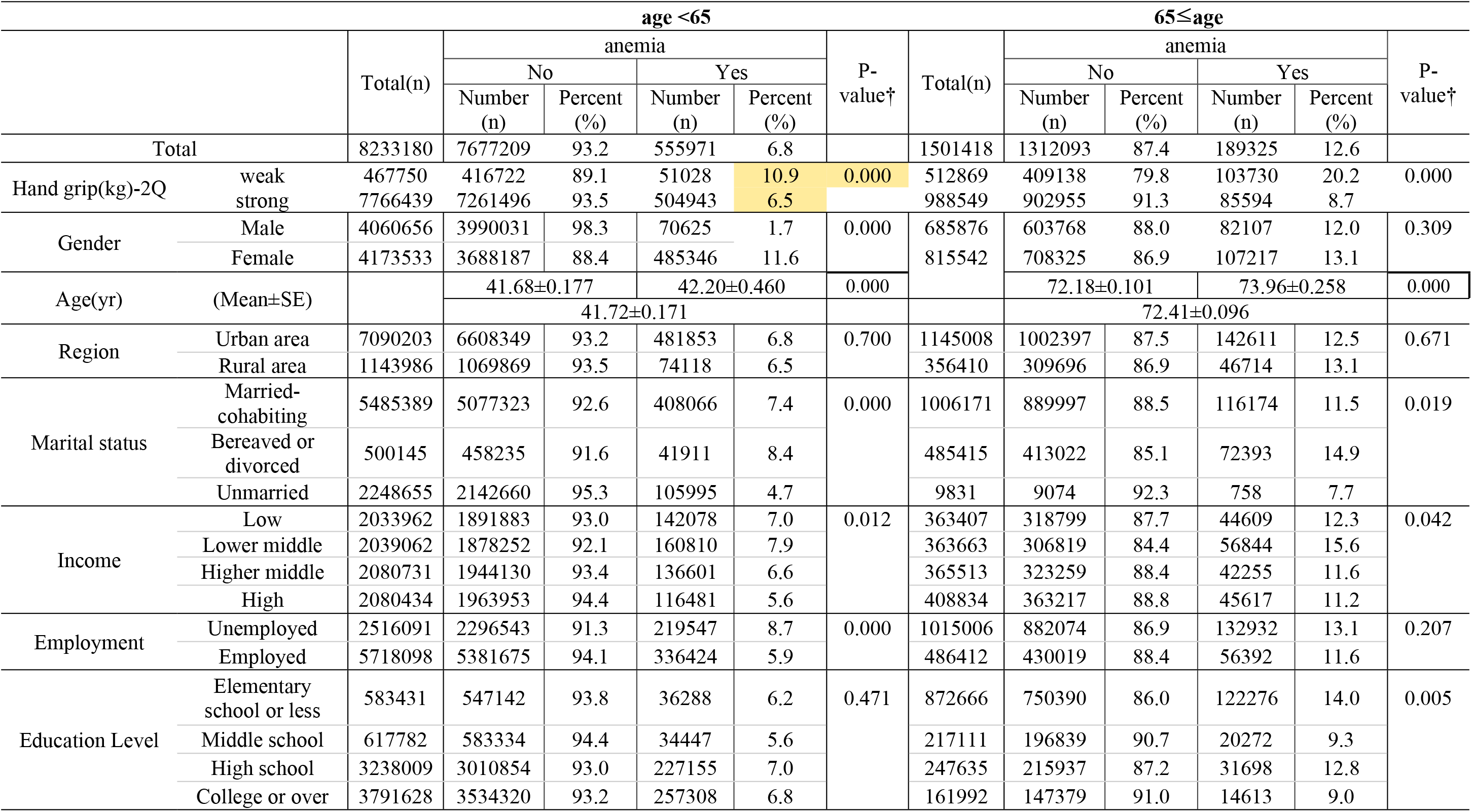
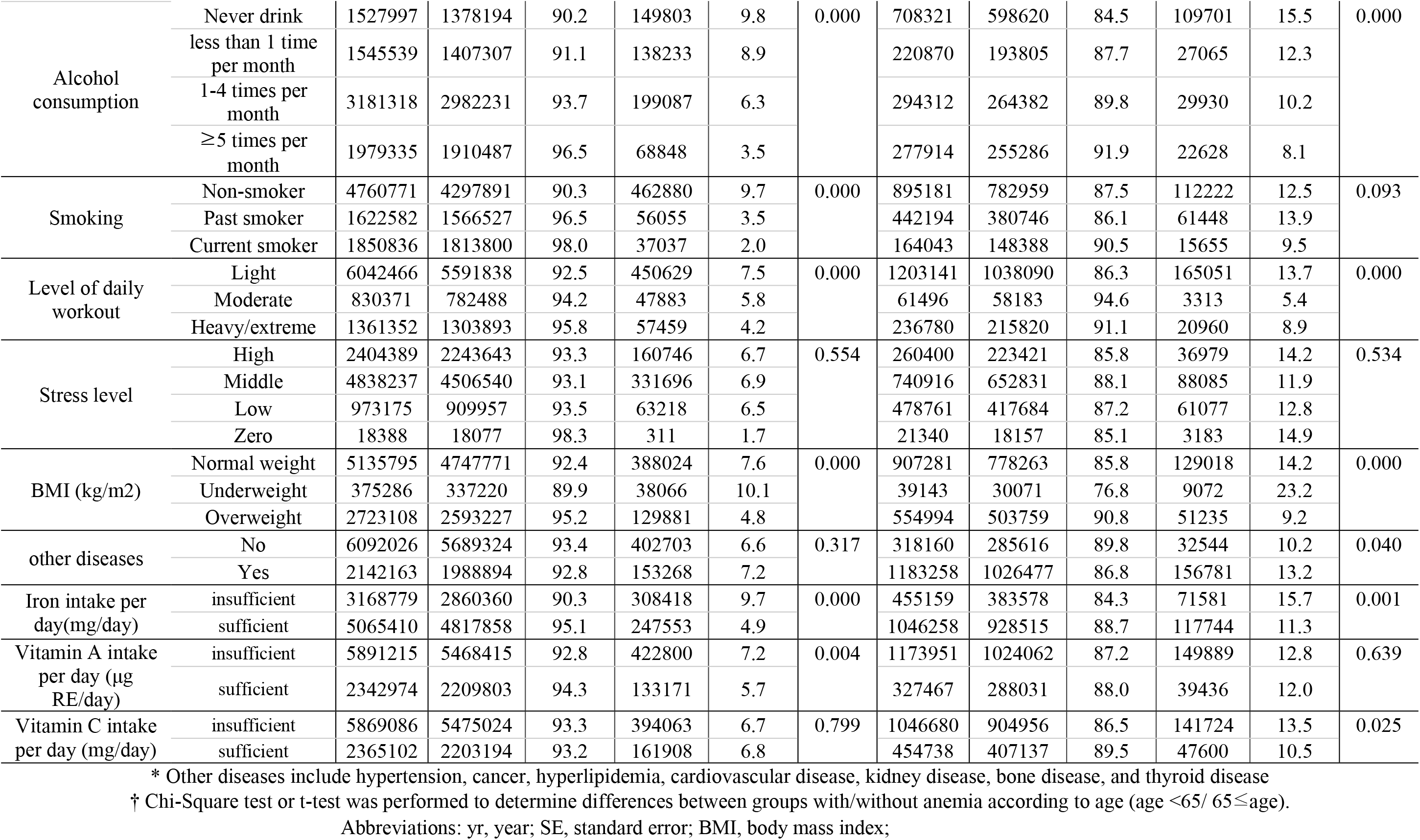
Characteristics of the study population by anemia according to age.

Table 4 shows the association between handgrip strength and anemia using logistic regression analyses. Model 1 adjusted for sociodemographic factors of anemia, such as sex, age, region, and income. Model 2 adjusted for individual lifestyle habit factors, such as smoking, alcohol consumption, and BMI, in addition to the sociodemographic factors. Finally, Model 3 shows association with anemia after adjusting for sociodemographic factors, individual lifestyle habit factors, and nutritional and disease-related factors.

**Table 4.**
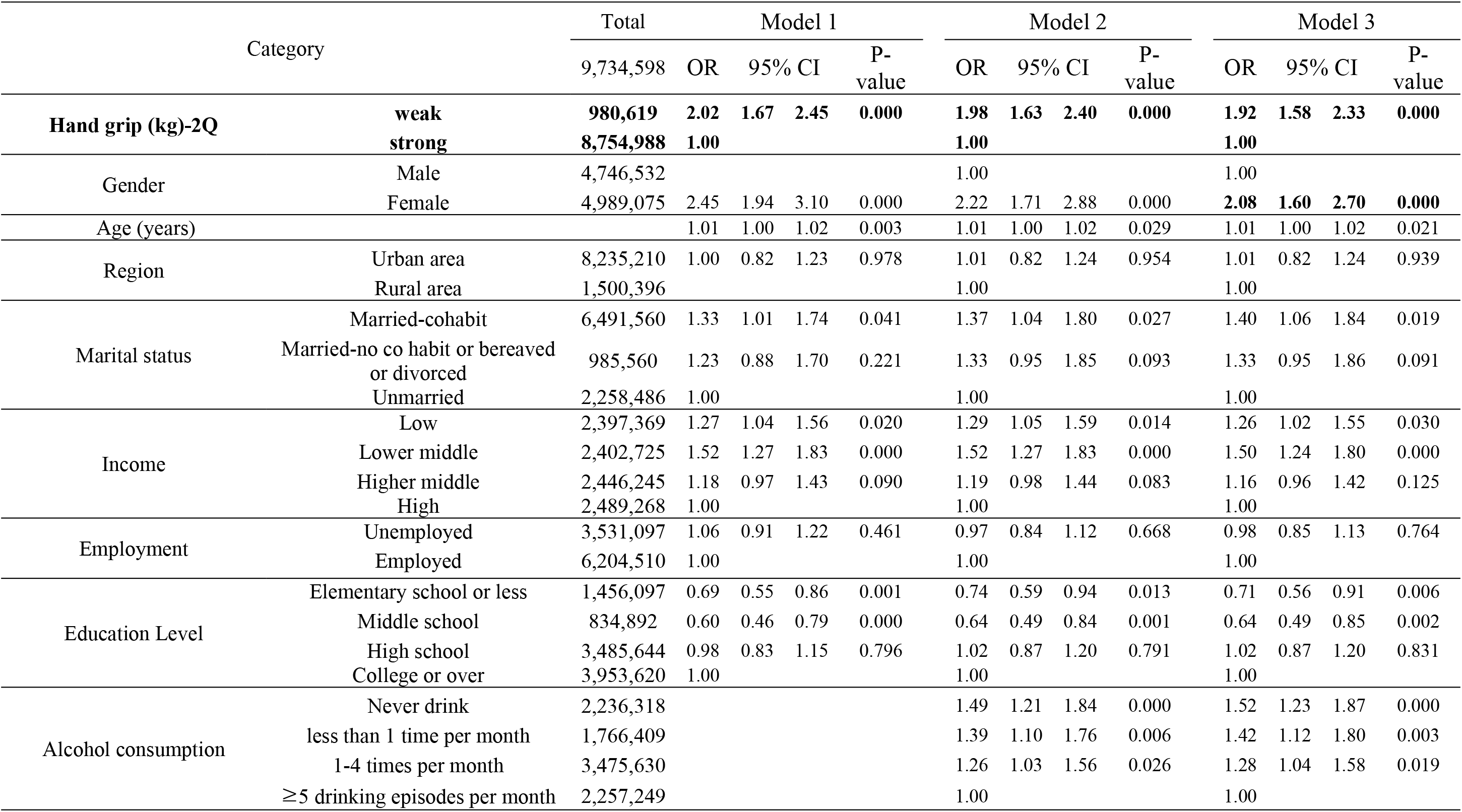
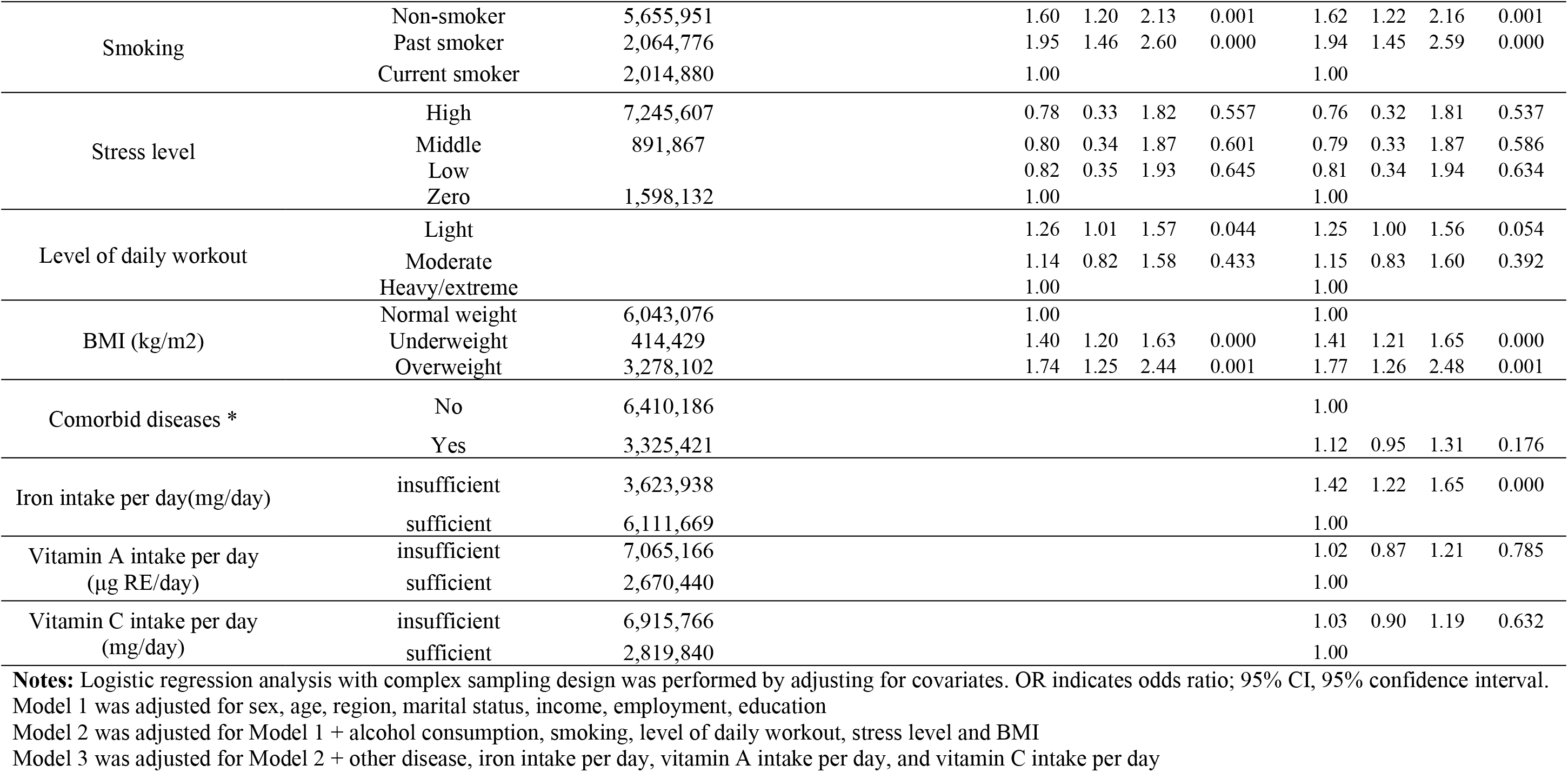
Association between hand grip and anemia.

The ORs of anemia for weak handgrip strength compared to the strong handgrip strength was 2.02, 1.98, and 1.92 in Models 1, 2, and 3, respectively, showing a significant association between weak handgrip strength and anemia (p<0.0001). The odds of having anemia were least for persons with 1-4 times per month frequency of alcohol consumption compared to the ≥5 drinking episodes per month persons, and highest in the light level of daily workout compared to the heavy/extreme groups. The odds ratio of underweight and overweight BMI was higher compared to the normal weight, and the odds of anemia was also higher in persons with comorbid diseases compared to persons without comorbid diseases at 1.12. Also, the prevalence of anemia was higher for persons with insufficient iron, vitamin A, and vitamin C intake per day compared to persons with sufficient intakes (Table 4).

Table 5 demonstrates the association between handgrip strength and anemia stratified by sex and age. As in Table 4, Model 1 shows the association with anemia while controlling for sociodemographic factors; Model 2 shows the values after controlling for individual lifestyle habit factors in addition to factors in Model 1; and Model 3 shows the values after controlling for sociodemographic factors, individual lifestyle habit factors, and nutritional and disease-related factors. Regardless of sex and age, there was a significant association between handgrip strength and anemia. In particular, the trend of weak handgrip strength in those with anemia compared to those without anemia was more pronounced in males than females (p=0.001, 95% CI: 1.35-3.34); and in those aged ≥65 years compared to those <65 (p<0.0001, 95% CI: 1.42-2.58).

**Table 5.** Association between hand grip and anemia by sub group.

| Class | Category |  | Model 1 |  |  |  | Model 2 |  |  |  | Model 3 |  |  |  |
| --- | --- | --- | --- | --- | --- | --- | --- | --- | --- | --- | --- | --- | --- | --- |
|  |  |  | OR | 95% CI |  | P-value | OR | 95% CI |  | P-value | OR | 95% CI |  | P-value |
| Male | Hand grip (kg)-2Q | weak | 2.36 | 1.53 | 3.63 | 0.000 | 2.11 | 1.34 | 3.31 | 0.001 | 2.13 | 1.35 | 3.34 | 0.001 |
|  |  | strong | 1.00 |  |  |  | 1.00 |  |  |  | 1.00 |  |  |  |
| Female | Hand grip (kg)-2Q | weak | 1.79 | 1.41 | 2.27 | 0.000 | 1.77 | 1.40 | 2.25 | 0.000 | 1.71 | 1.35 | 2.16 | 0.000 |
|  |  | strong | 1.00 |  |  |  | 1.00 |  |  |  | 1.00 |  |  |  |
| age <65 | Hand grip (kg)-2Q | weak | 1.59 | 1.18 | 2.15 | 0.003 | 1.58 | 1.17 | 2.14 | 0.003 | 1.57 | 1.16 | 2.12 | 0.003 |
|  |  | strong | 1.00 |  |  |  | 1.00 |  |  |  | 1.00 |  |  |  |
| 65≤ age | Hand grip (kg)-2Q | weak | 1.97 | 1.47 | 2.64 | 0.000 | 1.90 | 1.41 | 2.55 | 0.000 | 1.92 | 1.42 | 2.58 | 0.000 |
|  |  | strong | 1.00 |  |  |  | 1.00 |  |  |  | 1.00 |  |  |  |

Notes: Logistic regression analysis with a complex sampling design was performed by adjusting for covariates. OR indicate odds ratio; 95% CI, 95% confidence interval.
Model 1 was adjusted for sex, age, region, marital status, income, employment, education
Model 2 was adjusted for Model 1 + alcohol consumption, smoking, level of daily workout, stress level and BMI
Model 3 was adjusted for Model 2 + other disease, iron intake per day, vitamin A intake per day, and vitamin C intake per day

## Discussion

This study analyzed the association between handgrip strength and anemia in a total of 9,734,598 adults >19 years using a representative, reliable data that represents all Korean citizens. As a result, we found that handgrip strength and anemia had a clear correlation in the Korean adult population >19 years old. When handgrip strength was categorized into weak or strong based on the standards of the National Institutes of Health, the proportion of those having anemia with weaker hand grip strength was higher. Even after controlling for sociodemographic factors such as sex, age, region, and income, there was a clear correlation between handgrip strength and anemia. Furthermore, after controlling for individual lifestyle habit factors, such as smoking, alcohol consumption, nutritional and disease-related factors, handgrip strength and anemia showed a very significant association (OR=1.92, 95% CI: 1.58-2.33). Between the group with anemia and without anemia, indices such as BMI, level of daily workout, and iron intake per day showed significant differences. A peculiar finding is that the prevalence rate of anemia was higher for non-smokers and those with lower consumption of alcohol (Table 1). This is deduced to be due to the fact that females form a greater proportion of the anemia patients at 79.5% compared to males.

The relationship between anemia and handgrip strength can be explained by a biological mechanism as reported in previous studies. The low strength of anemia patients can be explained to be due to weakness and fatigue, and this can increase the risk of disability in these individuals [26]. Hemoglobin, in the red blood cells, is responsible for transporting oxygen to various parts of the body. Therefore, the lower blood concentration of hemoglobin means lower oxygen is transported, and this can increase the risk of various mobility disorders along with fatigue [27]. Because the hemoglobin concentration plays an important role in the oxygenation of the blood [18, 19], anemia defined by the level of hemoglobin represents reduction of oxygenation function. Hypoxia due to anemia can reduce the oxygen supply to the muscles and thereby affect the strength of the muscles [4].

We performed subgroup analyses by sex and age, which were predicted to affect anemia. In the subgroup by sex, the correlation between handgrip strength and anemia was higher for males compared to females. For males, indices such as BMI, level of daily workout, and comorbid disease had significant relationships with anemia (Table 2). Additionally, in the subgroups by age, the association between handgrip strength and anemia was more pronounced in subjects ≥65 years compared to those <65 years Although the proportion of females among anemia patients was far higher in the group <65 years, the male or female proportions did not show large differences in those ≥65 years. It was also found that BMI, level of daily workout, and iron intake per day were significantly related to anemia in both groups (Table 3).

Previous studies have shown significant relationships between anemia and handgrip strength in only females, rather than males. These include a study which found significant differences in hemoglobin levels in males among the South African population >40 years old [28] and a study which found an association between anemia and handgrip strength in Brazilian female population >60 years old [12]. However, this study showed higher odds of anemia in the weak handgrip strength group compared to the strong handgrip strength group in both males and females, and this association was greater in the male compared to the female group. The reason that the differences in the prevalence rates of anemia of males and females were different in each study can be the heterogeneity of the aging population with regards to race, life environment, and health problems, and these can all affect the hemoglobin levels [29].

Furthermore, previous studies on the elderly have found a clear association between handgrip strength and anemia. According to the research by Hirani V et al., the decline in hemoglobin concentration in the Australian elderly was directly related to the decline in handgrip strength [30]). Pennix et al also reported that anemia patients (defined by the hemoglobin level among elderly subjects >65 years), had far lower handgrip strength compared to patients without anemia [4]. Moreover, research on the association between anemia and handgrip strength in the elderly population >90 years old [28, 31] and research that found an association between anemia, handgrip strength, and lower-body strength in the elderly >100 years old [32] showed that the association between handgrip strength and association is large in the elderly population, especially as age increases. The reason could be that the reduction in handgrip strength due to aging can represent the overall health of the elderly [33]. In particular, because the elderly often have higher blood concentration of C-reactive protein, acute or chronic inflammation can result in an even larger physical decline [4, 5]. Therefore, the maintenance and management of handgrip strength are important especially in the elderly population, and it will be important to identify factors that affect handgrip strength [34].

There are several limitations with this research. First, because this study used a cross-sectional study design based on the KNHANES, caution must be taken to interpret a causal relationship between the variables. In other words, we cannot interpret as a causal relationship that handgrip strength is an independent causative factor of anemia or an epiphenomenon factor. Secondly, there is a limitation in identifying the risk factors of anemia because of insufficient nutritional data on folic acid or vitamin B12, which can cause anemia, since the effect of these nutritional vitamins on anemia could not be considered. It was difficult to clearly consider the effect of diseases that could be related to anemia. Thirdly, because KNHANES only considered Hb levels to define anemia, it was difficult to consider the types or causes of anemia. Therefore, additional research is necessary on various variables that can affect anemia; it is anticipated that the results of such a study can be used for clinical purposes. Despite these limitations, this research has the following advantages. First, this study is the first research to examine the correlation between handgrip strength and anemia in Korean adults. Although the fact that the concentration of hemoglobin can be directly related to handgrip strength has been found through previous studies, there was still insufficient research to confirm the association between anemia and handgrip strength. Because this research used data on the general population of Korea, the reliability of the data is high. Secondly, the results of this study showed that the relationship between handgrip strength and anemia was very strong; thus, it was suggested that handgrip strength can be used as a useful measurement tool for screening for anemia. Through this study, it will be possible to check and clinically follow up on the population with high likelihood of anemia through handgrip strength measurement, which is a simple tool that represents strength.

## Conclusion

The results of the present study showed that there was a strong association between handgrip strength and anemia. In particular, the association between handgrip strength and anemia was more pronounced for males and those who are older than 65. Therefore, handgrip strength can be used as a risk factor when screening for anemia in patients, and can be used in clinical practice for testing for anemia in the elderly, in epidemiological studies, or during treatments.

## Acknowledgements

### Disclosure of interests

The authors have declared that no competing interests exist.

### Author Contributions

Conceptualization, Resources: IHH BJ. Data curation: BJ. Formal analysis, Visualization, Writing (Original draft): YMG BJ. Methodology, Project administration, validation: KWK JHC. Software: BJ Supervision: IHH. Investigation, Writing (Review&Editing): YMG BYJ IHH

